# Interictal epileptiform discharges in focal epilepsy are preceded by a gradual increase in low-frequency oscillations

**DOI:** 10.1101/2020.05.27.118802

**Authors:** Karin Westin, Gerald Cooray, Daniel Lundqvist

## Abstract

Epilepsy is characterized by recurrent seizures and may also have negative influence on cognitive function. In addition to ictal activity, the epileptic brain also gives rise to *interictal epileptiform discharges* (IEDs). These IEDs constitute the diagnostic hallmark of epilepsy, and have been linked to impaired memory formation and negative effects on neurodevelopment. The neurophysiological dynamics underlying IED generation seem to resemble those underlying seizure development. Understanding the neurophysiological characteristics surrounding and preceding IED development would hence provide valuable insights into the pathophysiology of the epileptic brain. In order to improve this understanding, we aimed to characterize the dynamical activity changes that occurs immediately prior to an IED onset. We used magnetoencephalography (MEG) recordings from nine focal epilepsy patients to characterize the oscillatory activity preceding IED onsets. Our results showed a systematic and gradual increase in oscillatory delta and theta band activity (1-4 Hz and 4-8 Hz, respectively) during this pre-IED interval, reaching a maximum power at IED onset. These results indicate that the pre-IED brain state is characterized by a gradual synchronization that culminates in the neuronal hypersynchronization underlying IEDs. We discuss how IED generation might resemble seizure development, where physiological brain activity similarly undergoes a gradual synchronization that terminates in seizure onset.

## 1 Introduction

Epilepsy is one of the most common neurological disorders with prevalence of 4.3-7.8 per 1000 persons in developed countries, and with a total European yearly cost of disease close to EUR 15 billion (Pugliatti et al., 2007) Possible clinical manifestations include seizures, cognitive impairment and affected neurodevelopment. These symptoms can result from epileptic brain activities, including seizures and interictal epileptiform discharges (IEDs) (Badawy et al., 2012; Avanzini et al., 2013; Horak et al., 2017). However, although IEDs constitute one of the most important diagnostic hallmarks of epilepsy and can be used to localize epileptogenic foci (Asano et al., 2003; Gotman and Pittau, 2011) the relationship between the brain’s function during ictal and interictal phenomena remains inconclusive. Furthermore, IEDs appear able themselves to induce global network changes (Wilke et al., 2011; Malinowska et al., 2014; Lagarde et al., 2018), even mediating epilepsy-associated cognitive impairment and affecting normal neurodevelopment in childhood (Kobayashi et al., 2005, 2006; Fahoum et al., 2012; Ibrahim et al., 2014). For instance, large-scale physiological network communication is disrupted by IED-induced oscillations (Dahal et al., 2019), resulting in impaired memory formation (Gelinas et al., 2016; Tong et al., 2019).

In epilepsy, brain function can be considered a changeable state that can go from a normal state, to a state of IED generation, to a seizure state. An individual IED is considered to reflect a hyper-synchronized firing of a neuronal population (Jirsa et al., 2014; Jacob et al., 2019). In support of this view, several computational and animal models have demonstrated that IEDs and their cellular correlate, the paroxysmal depolarizing shift, can be triggered to occur *momentously* by increasing the excitation/inhibition ratio (Ayala et al., 1970; Schwartzkroin and Prince, 1979; Voskuyl and Albus, 1985; Mccormick and Contreras, 2001). However, analysis of human IED generation demonstrate that that there are additional changes in the brain state *prior* to IED onset, indicating that the context surrounding IED events develop over a longer time scale (Keller et al., 2010; Faizo et al., 2014; Bourel-Ponchel et al., 2017). Hemodynamical activation, for instance, peak one second before IEDs occur (Hawco et al., 2007; Jacobs et al., 2009; Rathakrishnan et al., 2010). Such changes are seen not only in the region generating IEDs – the so-called irritative zone (Jehi, 2018) – but also at distant regions, accompanied by altered connectivity patterns (Kobayashi et al., 2006; Faizo et al., 2014). Similarly, an HD-EEG study on benign childhood epilepsy with centrotemporal spikes (BECTS) demonstrate that the pre-IED state can be characterized both by increased synchronization and desynchronization (Bourel-Ponchel et al., 2017). Although it is unclear whether these pre-IED changes cause the IED generation, or whether they are mere reflections of the neural population activity resulting in IEDs, such changes further indicate that the neuronal activity build-up occurs slowly. In sum, the literature indicates that IEDs are not merely transient, momentous events confined to the irritative zone, but are generated in the context of slowly changing brain states that in turn influence the function of both epileptic and non-epileptic brain areas. However, such pre-IED brain state changes have yet to be fully characterized. Although pre-IED activity changes have been characterized in BECTS, this epilepsy diagnosis constitutes a specific, self-limited and age-related syndrome (Heijbel et al., 1975). Other epilepsies with different underlying etiologies (Manford, 2017) might exhibit different pre-IED changes. Furthermore, the temporal evolution of pre-IED brain state changes have not been characterized for any epilepsy diagnosis, and an increased understanding of this process might give an insight into the dynamical changes that mediate IED generation.

In this study, we aimed to analyze IED-related brain state fluctuations across focal epilepsies of different origin and patient age. More specifically, we aimed to *characterize the temporal characteristics of the pre-IED state*. For this purpose, we used magnetoencephalography (MEG) measurements from nine focal epilepsy patients undergoing MEG as part of non-invasive clinical epilepsy evaluation. Using these data, we explored changes in the oscillatory brain activity occurring prior to IED onset, and compared the characteristics of these periods to the oscillatory activity during periods free from IEDs.

## 2 Material and methods

### 2.1 Ethical approval

The experiment was approved by the Swedish Ethical Review Authority (DNR: 2018/1337-31), and was performed in agreement with the Declaration of Helsinki.

### 2.2 Patients

Nine patients diagnosed with pharmacoresistant focal epilepsy underwent a clinical MEG recording as part of epilepsy evaluation at the department of clinical neurophysiology at Karolinska University Hospital during 2017-2018 agreed to and were included in the study. Both children (n=2) and adults were included (median age = 33 years, 3 females). For demographics and focus localization, see Table 1.

**Table 1.** Patient demographics, number of IEDs, type and localization of IEDs.

| Demographics | IED source localization | No of IEDs | Type of IED |
| --- | --- | --- | --- |
| Patient 1 | Left frontal lobe | 27 | Spike and wave complex |
| Patient 2, | Right parietal/temporal lobe | 34 | Polyspikes |
| Patient 3 | Right medial orbitofrontal | 17 | Polyspike |
| Patient 4 | Left temporal | 14 | Spike/sharp wave |
| Patient 5 | Right temporal | 4 | Spike |
| Patient 6 | Right postcentral | 17 | Polyspike |
| Patient 7 | Frontal/temporal lobe, bilateral | 7 | Spike/sharp wave |
| Patient 8 | Temporal lobe, bilateral | 7 | Spike/Sharp wave |
| Patient 9 | Temporal lobe, bilateral | 7 | Spike/sharp wave |

### 2.3 MEG data acquisition and procedure

An Elekta Neuromag TRIUX 306-channel MEG system with 204 planar gradiometers and 102 magnetometers was used to record MEG data. Recordings were carried out at Karolinska Institutet (NatMEG facility) in a two-layer magnetically shielded room (Vacuumschmelze GmbH, model Ak3B). Data was sampled at 5000 Hz with an online 0.1 Hz high-pass filter and a 1650 Hz low-pass filter. Head-position indicator coils (HPI) were used to sample head movement during the recording. These were attached to the patients’ head and digitalized using Polhemus Fastrak motion tracker. During the recording, patients were positioned in supine position, with eyes closed and were asked to stay awake. The recording lasted approximately one hour.

### 2.4 Data preprocessing

Data pre-processing was performed using MaxFilter (Taulu and Simola, 2006) signal-space separation with buffer length 10 seconds and cut-off correlation coefficient 0.98. Raw data was bandpass filtered at 1-40 Hz using a Butterworth filter.

#### 2.4.1 IED source reconstruction

IEDs were located by visual inspection performed by an experienced physician (KW, the main author). Source reconstruction of interictal spikes/sharp waves and spike and slow wave complexes averaged across events was performed using Minimum Norm Estimate (MNE) (Hämäläinen and Ilmoniemi, 1994). Polyspike source reconstruction was performed using beamformer Dynamic Imaging of Coherent Sources (DICS) (Gross et al., 2001). The patients’ clinical MRI were utilized to create a full segmentation of the head and brain, which was performed using FreeSurfer (Dale et al., 1999; Fischl et al., 1999). The MNE-C software watershed algorithm (Gramfort et al., 2013) determined skin, skull and brain surface boundaries based upon this segmentation. A source space and a single compartment volume conductor model based upon the surface boundaries were also created using MNE-C.

### 2.5 Data analysis

Data preceding IED onset, defined as the first visible deflection from baseline, was analyzed alongside IED-free data. To this end, *pre-IED epochs* and *IED-free epochs* were created. One-second windows ending at IED onset and staring at least three seconds after any preceding IED constituted the pre-IED epochs. *IED-free epochs* were one-second windows taken from IED-free data at least three seconds away from any IED. The irritative zone was defined as the region to which the IEDs were located using source reconstruction. Pre-IED epochs were quantified for spectral and power composition in the irritative zone using four different methods described below. IED-free epochs were used for comparison. All analyses were performed using Python library MNE Python (Gramfort et al., 2013)

#### 2.5.1 Beamformer analysis

Oscillations in the pre-IED and IED-free epochs were analyzed using beamforming DICS (Gross et al., 2001; Hillebrand and Barnes, 2002). The number of IED-free epochs matched the number of pre-IED epochs. Frequency bands 1-4 Hz, 4-8 Hz, 8-13 Hz, 13-25 Hz corresponding to delta, theta, alpha and beta activity were analyzed.

#### 2.5.2 Event-related synchronization/desynchronization

In order to evaluate any neuronal synchronization or desynchronization of the irritative zone prior to IED formation, event-related synchronization/desynchronization (ERS/ERD) of the delta, theta, alpha and beta band of the pre-IED epochs were determined (Pfurtscheller and Lopes Da Silva, 1999). For each channel, a non-parametric cluster level paired t-test (Maris and Oostenveld, 2007) was performed to test whether any consistent power change occurred across epochs.

#### 2.5.3 Power spectral density (PSD) fluctuations

Source space data from the irritative zone located using source reconstruction as described above was extracted using MNE Python built-in functions (Gramfort et al., 2013). PSD of pre-IED and IED-free epochs was determined. The frequency band exhibiting pre-IED synchronization using DICS beamforming and/or event-related synchronization was analyzed. For both conditions, an array of maximum PSD of each epoch was created. For these two arrays, the minimum, maximum, mean, median and standard deviation were determined. The region contralateral to the irritative zone was analyzed in all patients.

#### 2.5.4 Temporal evolution of pre-IED frequency content

Averaged pre-IED irritative zone source space data was convoluted with sinus functions with frequencies 2-7 Hz, (one sinus function per integer frequency 2,3,4,5,6,7 Hz). Any frequency predominant in both time series will be enhanced in the resulting, convoluted time series, and all other frequencies will be damped. Amplitude of the resulting convoluted signal reflecting the temporal evolution of the neural time series frequency content was extracted as a function of time. A linear fit was performed in order to capture the temporal development of frequency-specific amplitude changes during pre-IED epochs.

## 3 Results

### 3.1 Source reconstruction

Patient demographics, number of IEDs, type and localization of IEDs can be found in Table 1.

### 3.2 Beamformer

Application of DICS beamformer revealed that six patients exhibited an enhanced theta band (4-8 Hz) and two patients exhibited an enhanced oscillatory delta band (1-4 Hz) activity at the irritative zone during pre-IED epochs compared to the rest of the cortical surface. In addition, a weaker oscillatory activity over the irritative zone was seen in four patients during the IED-free epochs compared to the rest of the cortical surface. See Figure 1 for two illustrative cases.

**Figure 1:**
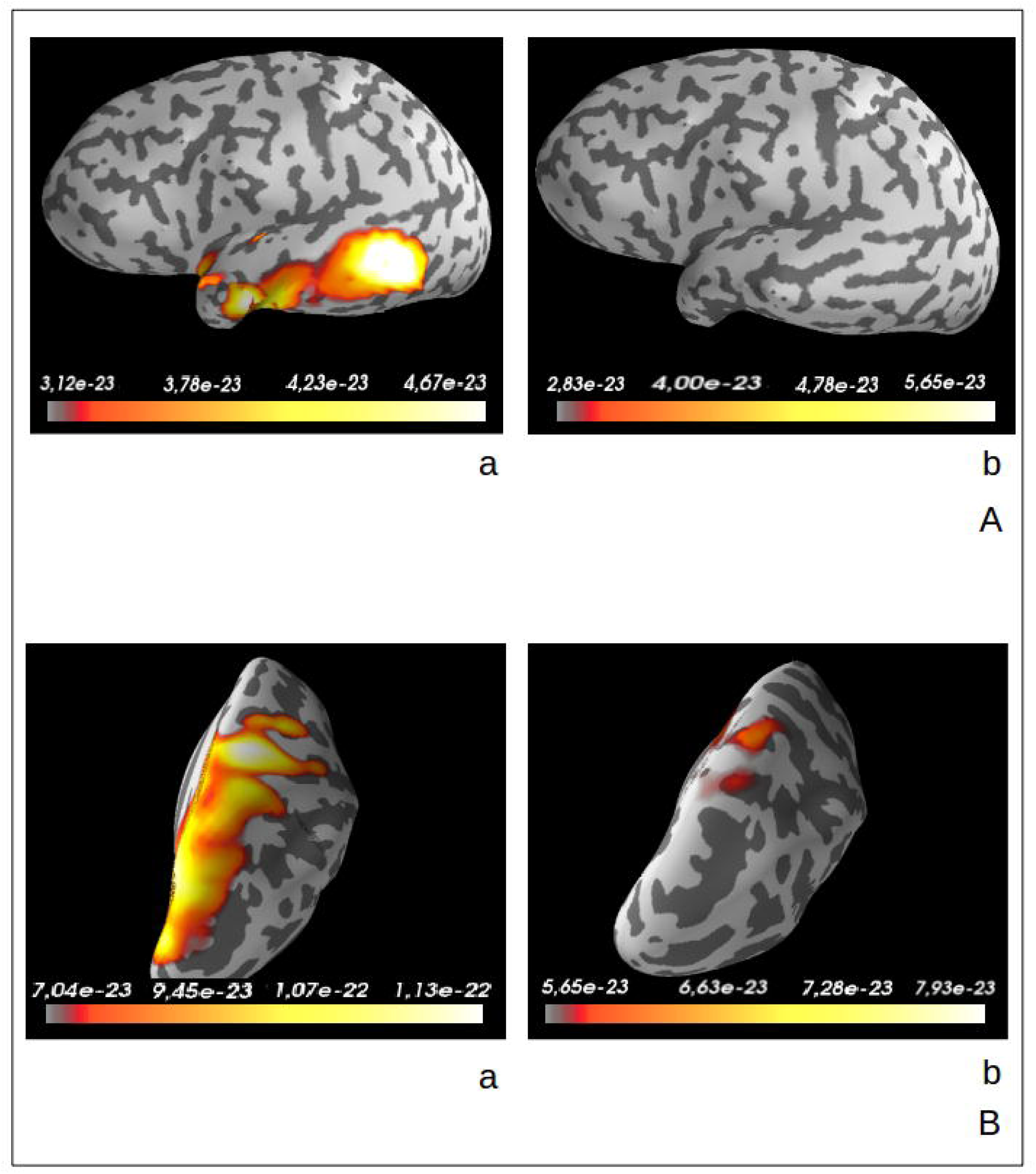
Two examples of theta oscillations located by DICS beamforming in patient 4 (Fig. 1A) and in patient 6 (Fig. 1B) during pre-IED epochs and during IED free epochs. Unit: Amperemeter

### 3.3 ERD-ERS

Six patients exhibited an increased synchronization within the theta or delta band in the pre-IED epochs in sensors covering the IED-generating zone. See Figure 2 for examples of such synchronization.

**Figure 2:**
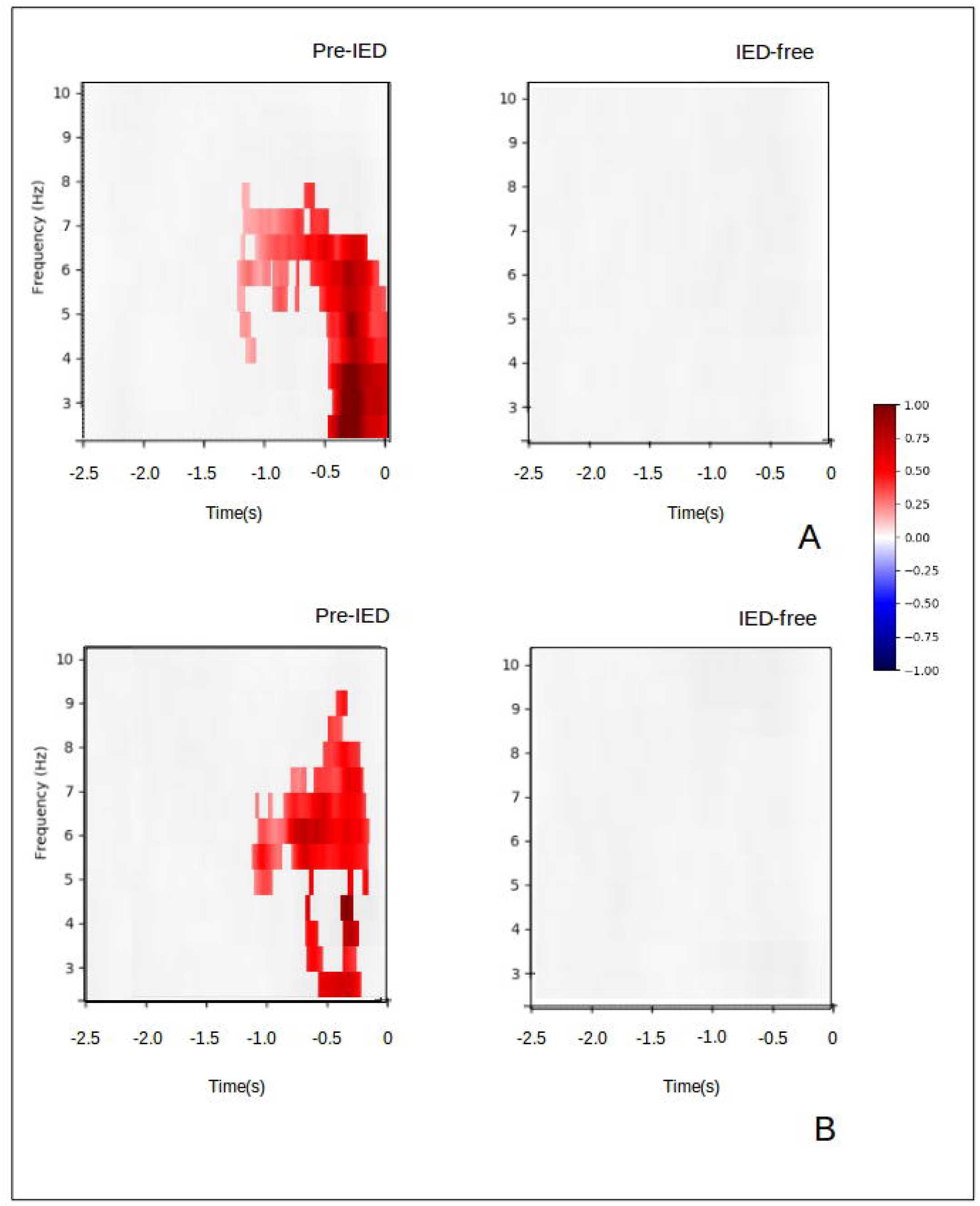
Two examples of event-related synchronization during pre-IED and IED-free epochs in two sensors covering the irritative zone in patient 1 (Fig 2A) and patient 3 (Fig 2B). The colorbar (red: synchronization, blue: desynchronization) indicates percent change in power (_μ_V^2^/Hz) compared to baseline (3 seconds preceding averaged pre-IED epochs).

### 3.4 PSD fluctuations

All but two patients exhibited a higher median PSD in pre-IED epochs compared to PSD in non-IED epochs. Of the six patients with unilateral epileptic foci, five exhibited the same pattern on the contralateral side. For all patients with unilateral foci, the median PSD levels of the pre-IED epochs were higher than those of the contralateral side. (See Table 2 and 3 for epileptogenic focus and the contralateral brain region in patients exhibiting unilateral epileptogenic foci; Table 4 and 5 for left and right hemisphere foci in patients exhibiting bilateral epileptogenic foci).

**Table 2:** PSD from the irritative zone in patients exhibiting unilateral epileptic foci

|  |  | Pre-IED PSD<br>(fT/cm<sup>2</sup>/Hz) | IED-free PSD<br>(fT/cm<sup>2</sup>/Hz) | Percentage increase pre-IED vs IED-free |
| --- | --- | --- | --- | --- |
|  |  | Median | Median | Median |

| Patient no | Frequency band |  |  |  |
| --- | --- | --- | --- | --- |
| Patient 1 | Theta | 993.6 | 826.2 | 20% |
| Patient 2 | Theta | 37.0 | 21.3 | 74% |
| Patient 3 | Theta | 3998.9 | 2843.3 | 41% |
| Patient 4 | Theta | 1084.6 | 819.9 | 32% |
| Patient 5 | Theta | 45.6 | 16.7 | 173% |
| Patient 6 | Theta | 13.5 | 6.4 | 111% |

**Table 3:** PSD from the brain region contralateral to the irritative zone in patients exhibiting unilateral epileptic foci

|  |  | Pre-IED PSD<br>(fT/cm<sup>2</sup>/Hz) | IED-free PSD<br>(fT/cm<sup>2</sup>/Hz) | Percentage increase pre-IED vs IED-free |
| --- | --- | --- | --- | --- |
|  |  | Median | Median | Median |
| Patient no | Frequency band |  |  |  |
| Patient 1 | Theta | 596.2 | 517.9 | 15% |
| Patient 2 | Theta | 20.0 | 12.5 | 60% |
| Patient 3 | Theta | 2746.5 | 2014.9 | 36% |
| Patient 4 | Theta | 31.0 | 32.1 | -3% |
| Patient 5 | Theta | 6.2 | 5.9 | 5% |
| Patient 6 | Theta | 7.3 | 5.5 | 33% |

**Table 4:** PSD from epileptic foci of the left hemisphere in patients exhibiting bilateral epileptic foci

|  |  | **Pre-IED PSD**<br>**(fT/cm<sup>2</sup>/Hz)** | **IED-free PSD**<br>**(fT/cm<sup>2</sup>/Hz)** | **Percentage increase**<br>**pre-IED vs IED-free** |
| --- | --- | --- | --- | --- |
|  |  | **Median** | **Median** | **Median** |
| **Patient no** | **Frequency band** |  |  |  |
| Patient 7 | Theta | 11.7 | 12.0 | -3% |
| Patient 8 | Delta | 4.0 | 1.3 | 208% |
| Patient 9 | Delta | 17.0 | 22.2 | -23% |

**Table 5:** PSD from epileptic foci of the right hemisphere in patients exhibiting bilateral epileptic foci

|  |  | **Pre-IED PSD**<br>**(fT/cm<sup>2</sup>/Hz)** | **IED-free PSD**<br>**(fT/cm<sup>2</sup>/Hz)** | **Percentage increase pre-**<br>**IED vs IED-free** |
| --- | --- | --- | --- | --- |
|  |  | **Median** | **Median** | **Median** |
| **Patient no** | **Frequency band** |  |  |  |
| Patient 7 | Theta | 11.7 | 12.0 | -3% |
| Patient 8 | Delta | 3.9 | 2.1 | 86% |
| Patient 9 | Delta | 3.5 | 4.6 | -24% |

### 3.5 Temporal evolution of pre-IED frequency content

In all but one patient, the power of oscillations within frequency band 4-6 Hz increased during the pre-IED epoch, reaching a maximum at IED onset. No such increase in power was seen outside of this interval.

## 4 Discussion

The aim of this study was to characterize the dynamics of oscillatory brain activity occurring prior to IED onset in the irritative zone. MEG measurements from nine focal epilepsy patients were used to characterize changes in oscillatory activity prior to IED onset. Such pre-IED epochs were compared to IED-free epochs. Our results show that IEDs were preceded by a power increase in low-frequency oscillations, thus demonstrating that the brain goes through a pre-IED state that is characterized by a low-frequency synchronization in the irritative zone (Timofeev and Steriade, 2004; Schnitzler and Gross, 2005). Furthermore, the power of these low-frequency oscillations grew throughout the pre-IED epoch, reaching its maximum at IED onset. This build-up indicates that the pre-IED state is characterized by a gradually increasing focal synchronization that culminates in an IED event. The result presented here thus indicate that IED generation resembles the gradual synchronization that has been observed during transition from normal brain activity to seizure activity (Lehnertz and Elger, 1995; Le Van Quyen et al., 2005; Mormann et al., 2000).

In mathematical models of *seizure generation*, the epileptic brain moves from a normal state to a seizure state that is characterized by sustained hypersynchronization (Wendling et al., 2005; Stefan and da Silva, 2013; Jirsa et al., 2014; Naze et al., 2015). Patient EEG data indicates that this transition is characterized by a long-lasting (up to hours) state of increased oscillatory activity (2-30 Hz) (D’Alessandro et al., 2003; Mormann et al., 2000; Le Van Quyen et al., 2005). Under these circumstances, brain activity perturbations such as fluctuating synaptic noise or changes in inhibition/excitation ratio can bring the epileptic brain into a state of increasing synchronization, eventually evolving into seizure activity (Cook et al., 2013; Jirsa et al., 2014; Wendling et al., 2005). Similarly, both mathematical and animal models indicate that corresponding dynamical brain network activity changes might underlie also IED generation (Jefferys and Haas, 1982; Hufnagel et al., 2000; Janszky et al., 2001; Wendling et al., 2002; Chaing et al., 2013; Jirsa et al., 2014; Naze et al., 2015; Uva et al., 2015). Furthermore, brain activity perturbations can also induce localized oscillations that spread throughout the cortical network, possibly mediating seizure development (Stacey et al., 2011). It is conceivable that the build-up in oscillatory activity seen in this study could be induced by such perturbations, and that this activity increases in power by gradually recruiting the irritative zone until a synchronization level sufficient to allow an IED onset is generated. In support of this view, our data showed that median power levels of delta and theta activity were increased during the pre-IED epochs compared to the IED-free epochs, suggesting that IEDs do not occur unless the irritative zone has reached a threshold level of synchronization. Similarly, microelectrode recordings from single neurons from the irritative zone demonstrate that a subregional increased neuronal firing occurs prior to IED onset (Keller et al., 2010). However, generation of an IED identifiable on scalp EEG recordings require simultaneous firing of at least 3 cm^2^ cortex (Tao et al., 2005), suggesting that the synchronized neuronal firing begins in small scale and then spread throughout the irritative zone until it reaches an extension where it can also be recorded with scalp EEG. The gradual increase in low-frequency power seen in our MEG recordings might correspond to such an increased involvement of the irritative zone. It is possible that the pre-IED state described here resembles the increased synchronization seen prior to seizure onset (Cámpora et al., 2019), indicating that generation of both IEDs and ictal activity might share common underlying neurophysiological dynamics.

### 4.1 Conclusion

Previous studies have shown that changes in synchronization of oscillatory brain activity occurs prior to IED onset (Keller et al., 2010; Faizo et al., 2014; Bourel-Ponchel et al., 2017). However, it has not been clear whether these pre-IED oscillations exhibit a dynamical evolution, with increasing synchronization in the irritative zone. In this study, we found evidence that IED onsets are preceded by a gradual power increase in low-frequency oscillations. These results resemble models of seizure development, where brain activity gradually becomes increasingly synchronized until an epileptic seizure is generated. It is noteworthy that pre-ictal changes are long-lasting, sometimes visible by visual inspection (Mormann et al., 2000; D’Alessandro et al., 2003; Le Van Quyen et al., 2005; Lehnertz et al., 2009), while the pre-IED changes reported here occur on a shorter time scale and requires averaging across epochs. It is conceivable that both IEDs and seizures arise from gradually increasing synchronization, but that the amount of synchronization determines whether a seizure or an IED occurs.

## Notes

### Competing Interest Statement

The authors have declared no competing interest.

## References

Asano, E., Muzik, O., Shah, A., Juhász, C., Chugani, D. C., Sood, S., et al. (2003). Quantitative interictal subdural EEG analyses in children with neocortical epilepsy. Epilepsia 44, 425–434. doi:10.1046/j.1528-1157.2003.38902.x.

Avanzini, G., Depaulis, A., Tassinari, A., and De Curtis, M. (2013). Do seizures and epileptic activity worsen epilepsy and deteriorate cognitive function? Epilepsia 54, 14–21. doi:10.1111/epi.12418.

Ayala, G. F., Matsumoto, H., and Gumnit, R. J. (1970). Excitability changes and inhibitory mechanisms in neocortical neurons during seizures. J. Neurophysiol. 33, 73–85. doi:10.1152/jn.1970.33.1.73.

Badawy, R. A. B., Johnson, K. A., Cook, M. J., and Harvey, A. S. (2012). A mechanistic appraisal of cognitive dysfunction in epilepsy. Neurosci. Biobehav. Rev. 36, 1885–1896. doi:10.1016/j.neubiorev.2012.05.002.

Bourel-Ponchel, E., Mahmoudzadeh, M., Berquin, P., and Wallois, F. (2017). Local and distant dysregulation of synchronization around interictal spikes in BECTS. Front. Neurosci. 11, 1–17. doi:10.3389/fnins.2017.00059.

Cámpora, N. E., Mininni, C. J., Kochen, S., and Lew, S. E. (2019). Seizure localization using pre ictal phase-amplitude coupling in intracranial electroencephalography. Sci. Rep. 9, 1–8. doi:10.1038/s41598-019-56548-y.

Chaing, C.-C., Lin, C.-C. K., Ju, M.-S., and Durand, D. M. (2013). High frequency stimulation can suppress globally seizures induced by 4-AP in the rat hippocampus: An acute in vivo stydy. Brain Stimlation 6, 180–189. doi:10.1038/jid.2014.371.

Cook, M. J., O’Brien, T. J., Berkovic, S. F., Murphy, M., Morokoff, A., Fabinyi, G., et al. (2013). Prediction of seizure likelihood with a long-term, implanted seizure advisory system in patients with drug-resistant epilepsy: A first-in-man study. Lancet Neurol. 12, 563–571. doi:10.1016/S1474-4422(13)70075-9.

D’Alessandro, M., Esteller, R., Echauz, J., Vachtsevanos, G., Hinson, A., and Litt, B. (2003). Epileptic Seizure Prediction Using Hybrid Feature Selection Over Multiple Intracranial EEG Electrode Contacts: A Report of Four Patients. IEEE Trans. Biomed. Eng. doi:10.1109/TBME.2003.810706.

Dahal, P., Ghani, N., Flinker, A., Dugan, P., Friedman, D., Doyle, W., et al. (2019). Interictal epileptiform discharges shape large-scale intercortical communication. Brain, 1–12. doi:10.1093/brain/awz269.

Dale, A. M., Fischl, B., and Sereno, M. I. (1999). Cortical Surface-Based Analysis I. Segmentation and Surface Reconstruction. Neuroimage 9, 179–194. doi:10.1006/nimg.1998.0395.

Fahoum, F., Lopes, R., Pittau, F., Dubeau, F., and Gotman, J. (2012). Widespread epileptic networks in focal epilepsies: EEG-fMRI study. Epilepsia 53, 1618–1627. doi:10.1111/j.1528-1167.2012.03533.x.

Faizo, N. L., Burianová, H., Gray, M., Hocking, J., Galloway, G., and Reutens, D. (2014). Identification of pre-spike network in patients with mesial temporal lobe epilepsy. Front. Neurol. 5, 1–8. doi:10.3389/fneur.2014.00222.

Fischl, B., Sereno, M. I., and Dale, A. M. (1999). Cortical surface-based analysis: II. Inflation, flattening, and a surface-based coordinate system. Neuroimage 9, 195–207. doi:10.1006/nimg.1998.0396.

Gelinas, J. N., Khodagholy, D., Thesen, T., Devinsky, O., and Buzsáki, G. (2016). Interictal epileptiform discharges induce hippocampal-cortical coupling in temporal lobe epilepsy. Nat. Med. 22, 641–648. doi:10.1016/j.physbeh.2017.03.040.

Gotman, J., and Pittau, F. (2011). Combining EEG and fMRI in the study of epileptic discharges. Epilepsia 52, 38–42. doi:10.1111/j.1528-1167.2011.03151.x.

Gramfort, A., Luessi, M., Larson, E., Engemann, D. A., Strohmeier, D., Brodbeck, C., et al. (2013). MEG and EEG data analysis with MNE-Python. Front. Neurosci. 7, 1–13. doi:10.3389/fnins.2013.00267.

Gross, J., Kujala, J., Hamalainen, M., Timmermann, L., Schnitzler, A., and Salmelin, R. (2001). Dynamic imaging of coherent sources: Studying neural interactions in the human brain. Proc. Natl. Acad. Sci. doi:10.1073/pnas.98.2.694.

Hämäläinen, M. S., and Ilmoniemi, R. J. (1994). Interpreting magnetic fields of the brain: minimum norm estimates. Med. Biol. Eng. Comput. 32, 35–42. doi:https://doi.org/10.1007/BF02512476.

Hawco, C. S., Bagshaw, A. P., Lu, Y., Dubeau, F., and Gotman, J. (2007). BOLD changes occur prior to epileptic spikes seen on scalp EEG. Neuroimage 35, 1450–1458. doi:10.1016/j.neuroimage.2006.12.042.

Heijbel, J., Blom, S., and Bergfors, P. G. (1975). Benign Epilepsy of Children with Centrotemporal EEG Foci. A Study of Incidence Rate in Outpatient Care. Epilepsia 16, 657–664. doi:10.1111/j.1528-1157.1975.tb04748.x.

Hillebrand, A., and Barnes, G. R. (2002). A quantitative assessment of the sensitivity of whole-head MEG to activity in the adult human cortex. Neuroimage 16, 638–650. doi:10.1006/nimg.2002.1102.

Horak, P. C., Meisenhelter, S., Song, Y., Testorf, M. E., Kahana, M. J., Viles, W. D., et al. (2017). Interictal epileptiform discharges impair word recall in multiple brain areas. Epilepsia 58, 373–380. doi:10.1111/epi.13633.

Hufnagel, A., Dümpelmann, M., Zentner, J., Schijns, O., and Elger, C. E. (2000). Clinical relevance of quantified intracranial interictal spike activity in presurgical evaluation of epilepsy. Epilepsia 41, 467–478. doi:10.1111/j.1528-1157.2000.tb00191.x.

Ibrahim, G. M., Cassel, D., Morgan, B. R., Smith, M. Lou, Otsubo, H., Ochi, A., et al. (2014). Resilience of developing brain networks to interictal epileptiform discharges is associated with cognitive outcome. Brain 137, 2690–2702. doi:10.1093/brain/awu214.

Jacob, T., Lillis, K. P., Wang, Z., Swiercz, W., Rahmati, N., and Staley, K. J. (2019). A proposed mechanism for spontaneous transitions between interictal and ictal activity. J. Neurosci. 39, 557–575. doi:10.1523/JNEUROSCI.0719-17.2018.

Jacobs, J., LeVan, P., Moeller, F., Boor, R., Stephani, U., Gotman, J., et al. (2009). Hemodynamic changes preceding the interictal EEG spike in patients with focal epilepsy investigated using simultaneous EEG-fMRI. Neuroimage 45, 1220–1231. doi:10.1016/j.neuroimage.2009.01.014.

Janszky, J., Fogarasi, A., Jokeit, H., Schulz, R., Hoppe, M., and Ebner, A. (2001). Spatiotemporal relationship between seizure activity and interictal spikes in temporal lobe epilepsy. Epilepsy Res. 47, 179–188. doi:10.1016/S0920-1211(01)00307-2.

Jefferys, J. G. R., and Haas, H. L. (1982). Synchronized bursting of CA1 hippocampal pyramidal cells in the absence of synaptic transmission. Nature 300, 448–450. doi:10.1038/300448a0.

Jehi, L. (2018). The epileptogenic zone: Concept and definition. Epilepsy Curr. 18, 12–16. doi:10.5698/1535-7597.18.1.12.

Jirsa, V. K., Stacey, W. C., Quilichini, P. P., Ivanov, A. I., and Bernard, C. (2014). On the nature of seizure dynamics. Brain 137, 2210–2230. doi:10.1093/brain/awu133.

Keller, C. J., Truccolo, W., Gale, J. T., Eskandar, E., Thesen, T., Carlson, C., et al. (2010). Heterogeneous neuronal firing patterns during interictal epileptiform discharges in the human cortex. Brain 133, 1668–1681. doi:10.1093/brain/awq112.

Kobayashi, E., Bagshaw, A. P., Grova, C., Dubeau, F., and Gotman, J. (2006). Negative BOLD responses to epileptic spikes. Hum. Brain Mapp. 27, 488–497. doi:10.1002/hbm.20193.

Kobayashi, K., Yoshinaga, H., Ohtsuka, Y., and Gotman, J. (2005). Dipole modeling of epileptic spikes can be accurate or misleading. Epilepsia 46, 397–408. doi:10.1111/j.0013-9580.2005.31404.x.

Lagarde, S., Roehri, N., Lambert, I., Trebuchon, A., McGonigal, A., Carron, R., et al. (2018). Interictal stereotactic-EEG functional connectivity in refractory focal epilepsies. Brain 141, 2966–2980. doi:10.1093/brain/awy214.

Le Van Quyen, M., Soss, J., Navarro, V., Robertson, R., Chavez, M., Baulac, M., et al. (2005). Preictal state identification by synchronization changes in long-term intracranial EEG recordings. Clin. Neurophysiol. 116, 559–568. doi:10.1016/j.clinph.2004.10.014.

Lehnertz, K., Bialonski, S., Horstmann, M. T., Krug, D., Rothkegel, A., Staniek, M., et al. (2009). Synchronization phenomena in human epileptic brain networks. J. Neurosci. Methods 183, 42–48. doi:10.1016/j.jneumeth.2009.05.015.

Lehnertz, K., and Elger, C. E. (1995). Spatio-temporal dynamics of the primary epileptogenic area in temporal lobe epilepsy characterized by neuronal complexity loss. Electroencephalogr. Clin. Neurophysiol. 95, 108–117. doi:10.1016/0013-4694(95)00071-6.

Malinowska, U., Badier, J. M., Gavaret, M., Bartolomei, F., Chauvel, P., and Bénar, C. G. (2014). Interictal networks in Magnetoencephalography. Hum. Brain Mapp. 35, 2789–2805. doi:10.1002/hbm.22367.

Manford, M. (2017). Recent advances in epilepsy. J. Neurol. 264, 1811–1824. doi:10.1007/s00415-017-8394-2.

Maris, E., and Oostenveld, R. (2007). Nonparametric statistical testing of EEG- and MEG-data. J. Neurosci. Methods 164, 177–190. doi:10.1016/j.jneumeth.2007.03.024.

Mccormick, D. A., and Contreras, D. (2001). On the cellular and network bases of epileptic seizures. 107, 96–107.

Mormann, F., Lehnertz, K., David, P., and E. Elger C. (2000). Mean phase coherence as a measure for phase synchronization and its application to the EEG of epilepsy patients. Phys. D Nonlinear Phenom. 144, 358–369. doi:10.1016/S0167-2789(00)00087-7.

Naze, S., Bernard, C., and Jirsa, V. (2015). Computational Modeling of Seizure Dynamics Using Coupled Neuronal Networks: Factors Shaping Epileptiform Activity. PLoS Comput. Biol. 11, 1–21. doi:10.1371/journal.pcbi.1004209.

Pfurtscheller, G., and Lopes Da Silva, F. H. (1999). Event-related EEG/MEG synchronization and desynchronization: basic principles. Clin Neurophsiol 110, 1842–1857. doi: 10.1016/s1388-2457(99)00141-8

Pugliatti, M., Beghi, E., Forsgren, L., Ekman, M., and Sobocki, P. (2007). Estimating the cost of epilepsy in Europe: A review with economic modeling. Epilepsia 48, 2224–2233. doi:10.1111/j.1528-1167.2007.01251.x.

Rathakrishnan, R., Moeller, F., Levan, P., Dubeau, F., and Gotman, J. (2010). BOLD signal changes preceding negative responses in EEG-fMRI in patients with focal epilepsy. Epilepsia 51, 1837–1845. doi:10.1111/j.1528-1167.2010.02643.x.

Schnitzler, A., and Gross, J. (2005). Normal and pathological oscillatory communication in the brain. Nat. Rev. Neurosci. 6, 285–296. doi:10.1038/nrn1650.

Schwartzkroin, P. A., and Prince, D. A. (1979). Changes in excitatory and inhibitory synaptic potentials leading to epileptogenic activity. Brain Res. 183, 61–77. doi:10.1016/0006-8993(80)90119-5.

Stacey, W. C., Krieger, A., and Litt, B. (2011). Network recruitment to coherent oscillations in a hippocampal computer model. J. Neurophysiol. 105, 1464–1481. doi:10.1152/jn.00643.2010.

Stefan, H., and da Silva, F. H. L. (2013). Epileptic neuronal networks: Methods of identification and clinical relevance. Front. Neurol. 4 MAR, 1–15. doi:10.3389/fneur.2013.00008.

Tao, J. X., Ray, A., Hawes-Ebersole, S., and Ebersole, J. S. (2005). Intracranial EEG Substrates of Scalp EEG Interictal Spikes. Epilepsia 45, 669–676. doi:10.1111/j.1528-1167.2005.11404.x.

Taulu, S., and Simola, J. (2006). Spatiotemporal signal space separation method for rejecting nearby interference in MEG measurements. Phys. Med. Biol. 51, 1759–1768. doi:10.1088/0031-9155/51/7/008.

Timofeev, I., and Steriade, M. (2004). Neocortical seizures: Initiation, development and cessation. Neuroscience 123, 299–336. doi:10.1016/j.neuroscience.2003.08.051.

Tong, X., An, D., Xiao, F., Lei, D., Niu, R., Li, W., et al. (2019). Real-time effects of interictal spikes on hippocampus and amygdala functional connectivity in unilateral temporal lobe epilepsy: An EEG-fMRI study. Epilepsia 60, 246–254. doi:10.1111/epi.14646.

Uva, L., Breschi, G. L., Gnatkovsky, V., Taverna, S., and de Curtis, M. (2015). Synchronous Inhibitory Potentials Precede Seizure-Like Events in Acute Models of Focal Limbic Seizures. J. Neurosci. 35, 3048–3055. doi:10.1523/jneurosci.3692-14.2015.

Voskuyl, R. A., and Albus, H. (1985). Spontaneous epileptiform discharges in hippocampal slices induced by 4-aminopyridine. Brain Res. 342, 54–66. doi:10.1016/0006-8993(85)91352-6.

Wendling, F., Bartolomei, F., Bellanger, J. J., and Chauvel, P. (2002). Epileptic fast activity can be explained by a model of impaired GABAergic dendritic inhibition. Eur. J. Neurosci. 15, 1499–1508. doi:10.1046/j.1460-9568.2002.01985.x.

Wendling, F., Hernandez, A., Bellanger, J.-J., Chauvel, P., and Bartolomei, F. (2005). Interictal to ictal transition in human temporal lobe epilepsy: insights from a computational model of intracerebral EEG. J. Clin. Neurophysiol. 22, 343–56. doi: 10.1016/s1388-2457(99)00141-8.

Wilke, C., Worrell, G. A., and He, B. (2011). Graph analysis of epileptogenic networks in human partial epilepsy. Epilepsia 52, 84–93. doi:10.1038/mp.2011.182.doi.

